# NAADP elicits two-pore channel currents by lifting Lsm12-mediated inhibition of PI(3,5)P_2_ activation

**DOI:** 10.64898/2026.04.13.718294

**Authors:** Xin Guan, Canwei Du, Kunal R. Shah, Jiusheng Yan

**Author notes:** Correspondence should be addressed to J.Y. These authors contributed equally (Xin Guan, Canwei Du, Kunal R. Shah).

## Abstract

Mammalian two-pore channels (TPCs) are endolysosomal cation channels that regulate membrane trafficking and ionic homeostasis and have been strongly implicated in nicotinic acid adenine dinucleotide phosphate (NAADP) signaling. We previously identified Lsm12 as an NAADP receptor and TPC-interacting protein required for NAADP-evoked Ca^2+^ mobilization from acidic organelles; however, how NAADP–Lsm12 coupling regulates TPC gating has remained unclear. Here, we show that Lsm12 acts as a potent antagonist of phosphatidylinositol 3,5-bisphosphate [PI(3,5)P_2_]–dependent TPC activation. Purified Lsm12 strongly inhibited PI(3,5)P_2_-evoked currents of TPC1 and TPC2, and endogenous Lsm12 similarly suppressed TPC2 activity in cells. Mechanistically, Lsm12 reduces the apparent sensitivity of TPC2 to PI(3,5)P_2_ through a competitive mechanism that depends on the concentrations of both Lsm12 and PI(3,5)P_2_, as well as on Lsm12–TPC interaction. Importantly, NAADP specifically and dose-dependently reverses Lsm12-mediated inhibition, restoring TPC currents only in the presence of PI(3,5)P_2_ or an intact PI(3,5)P_2_-binding site on TPCs. Consistently, acute sequestration of endogenous PI(3,5)P_2_ reduces NAADP-evoked cytosolic Ca^2+^ signals. These findings support a model in which Lsm12 tonically restrains PI(3,5)P_2_-dependent TPC gating, whereas NAADP binding to Lsm12 relieves this inhibition to permit channel activation. Our study therefore establishes a mechanistic link between NAADP signaling and phosphoinositide-dependent TPC gating and provides a working model for understanding NAADP-evoked Ca^2+^ release from acidic stores.

## INTRODUCTION

Two-pore channels (TPCs) are predominantly localized to acidic organelles of the endolysosomal system in animals and to vacuoles in plants. TPCs are homodimeric cation channels, with each subunit containing two homologous domains, each consisting of six transmembrane segments and a pore loop, resembling the basic structural unit of voltage-gated ion channels. In mammals, In mammals, two functional isoforms are present: TPC1, which is broadly distributed across endosomal and lysosomal compartments, and TPC2, which is primarily localized to late endosomes and lysosomes (*1, 2*). Mammalian TPCs are potently activated by phosphatidylinositol 3,5-bisphosphate (PI(3,5)P_2_) (*1–4*), inhibited by ATP via mTORC1 signaling (*5*). TPC1 exhibits voltage dependence (*6*), whereas TPC2 is largely insensitive to voltage (*3*). The reported structures of mouse TPC1 and human TPC2 have provided important insights into TPC architecture and gating by voltage and PI(3,5)P_2_ (*7, 8*). Functionally, endolysosomal TPCs regulate endomembrane dynamics and Ca²⁺ homeostasis within acidic organelles (*1, 9*). Accordingly, TPCs have been implicated in processes such as autophagy (*10*) and viral entry, including that of Ebola virus (*11*) and coronaviruses (*12, 13*). In addition, genetic and experimental studies have linked TPC2 to pigmentation (*14–16*) and metabolic (*17, 18*) and neurodegenerative phenotypes (*19*).

Accumulating evidence supports a critical role for TPCs in NAADP-evoked Ca²⁺ release from acidic stores (*20–26*). NAADP is a potent Ca²⁺-mobilizing second messenger and uniquely targets Ca²⁺ release from acidic endolysosomal compartments, in contrast to other second messengers that primarily act on the endoplasmic reticulum (*27–31*). This distinctive signaling pathway positions NAADP as a key regulator of localized Ca²⁺ dynamics associated with endolysosomal function. However, NAADP does not bind directly to TPCs (*32, 33*). We identified Lsm12, a Sm-like (Lsm) protein, as an NAADP receptor and a regulator of TPC function (*33*). Lsm12 interacts with TPCs and binds NAADP with high affinity and specificity relative to NADP. Genetic ablation of Lsm12 abolishes NAADP-evoked Ca²⁺ release, whereas re-expression or acute reintroduction of purified Lsm12 protein restores this response. Another protein, JPT2, has also been identified as an NAADP-binding protein (*34, 35*).

Despite these advances, several key questions remain unresolved. In particular, the mechanism by which NAADP activates TPCs via its receptor proteins remains unknown. PI(3,5)P_2_ is currently the only established endogenous direct activator of TPCs. It is unclear whether and how Lsm12, as a TPC-interacting protein, modulates PI(3,5)P_2_-dependent TPC gating. Furthermore, whether NAADP can evoke Ca²⁺-permeable currents through mammalian TPCs remains debated, given the relatively low Ca²⁺ permeability of PI(3,5)P_2_-activated TPCs (*1*) and the distinct structural features of their selectivity filters compared with those of Ca²⁺-permeable plant TPCs (*2*). NAADP-evoked TPC currents have been reported (*3, 22, 36–39*); however, the mechanism underlying NAADP-dependent activation of TPCs remains unresolved and robust currents have been difficult to detect in many patch-clamp recordings (*1, 2, 4–6*) and. These observations suggest that additional factors may be required to couple TPCs to NAADP signaling.

Here, we investigate whether and how Lsm12 and NAADP regulate TPC activity. We show that Lsm12 inhibits PI(3,5)P_2_-dependent TPC activation and that NAADP relieves this inhibition, thereby enabling TPC activation in the presence of PI(3,5)P_2_. Our findings thus establish a Lsm12-mediated functional link between NAADP signaling and PI(3,5)P_2_-dependent TPC activation and provide a potential framework for understanding NAADP-evoked Ca²⁺ release from acidic stores.

## RESULTS

### Purified Lsm12 antagonizes endolysosomal TPCs activated by PI(3,5)P_2_

We have previously reported that purified recombinant human Lsm12 is effective in NAADP binding and rapidly rescues NAADP-evoked Ca2+ signals upon its micro-injection into Lsm12 knockout (KO) HEK293 cells (*33*). Given that Lsm12 interacts with TPCs, we investigated how the purified Lsm12 affects TPC gating properties, particularly their activation by the endogenous ligand PI(3,5)P_2_. Using HEK293 cells transfected with TPC expression plasmids, we performed patch-clamp recordings of endolysosomes enlarged by vacuolin-1 and freshly microdissected from cells immediately before recording. In the presence of 0.3 µM PI(3,5)P_2_, we observed that purified Lsm12 (**Fig. 1A**), when perfused onto enlarged lysosomes at submicromolar and micromolar concentrations, significantly inhibited TPC2 currents in a dose-dependent manner (**Fig. 1B**). Lsm12 at 0.1, 0.5, 1, 2, and 5 µM inhibited PI(3,5)P_2_-activated TPC2 currents by 52%, 70%, 82%, 90%, and 95%, respectively. Similarly, purified Lsm12 at 2 µM resulted in a significant 78% inhibition of TPC1 currents activated by 0.3 µM PI(3,5)P_2_ (**Fig. 1C**). To evaluate whether Lsm12 inhibition of PI(3,5)P_2_-activated channels is specific to TPCs, we measured its effect on another lysosomal PI(3,5)P_2_-activated channel, TRPML1. We found that Lsm12 at 2 µM had no major effect on TRPML1 currents elicited by 0.3 µM PI(3,5)P_2_ (**Fig. 1D**).

**Figure 1.**
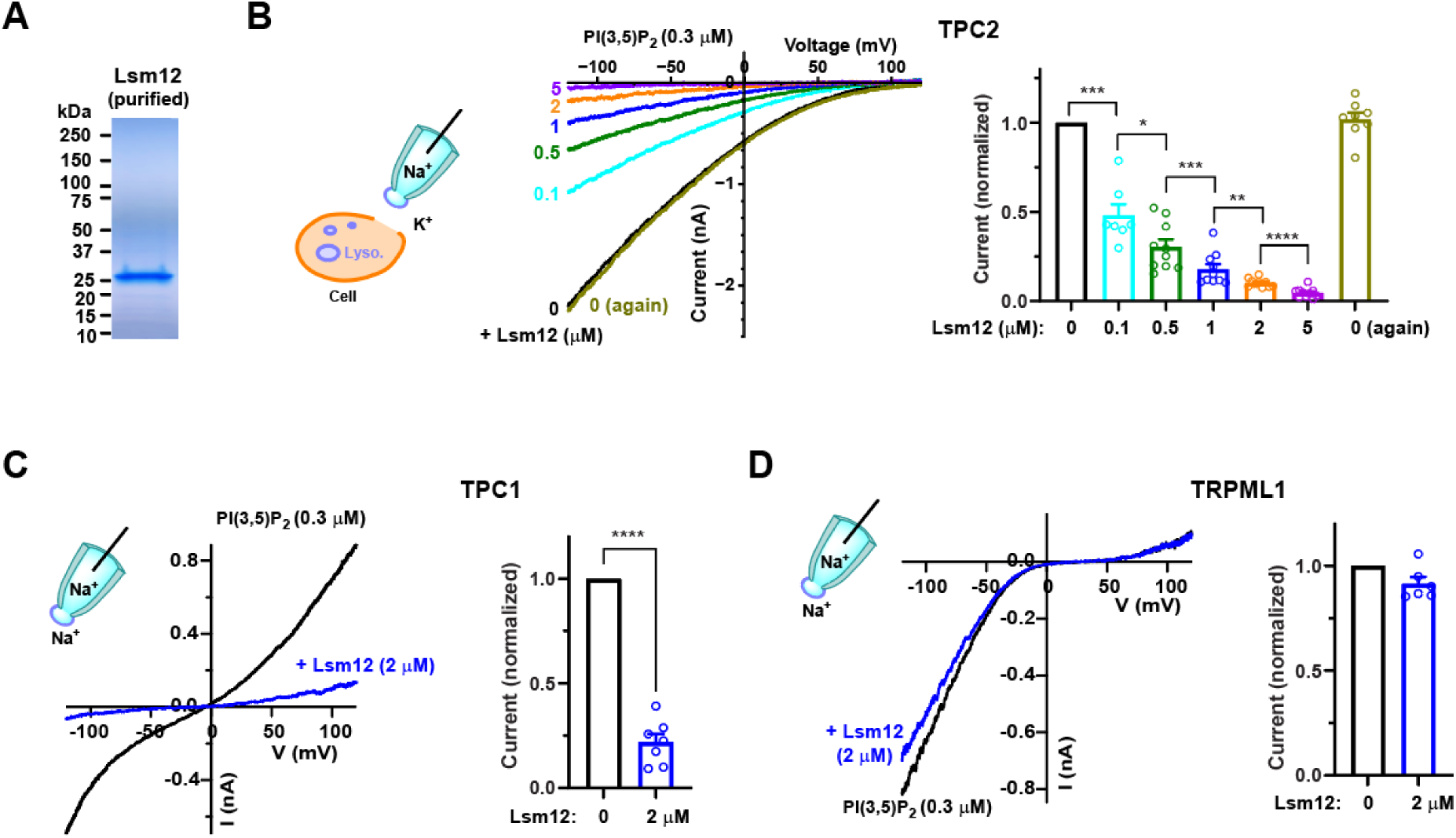
Reduction of endolysosomal TPC currents by purified Lsm12. **(A)** Purified Lsm12 protein used throughout this study. Coomassie blue staining of an SDS-PAGE gel shows a single band near 25 kDa, consistent with the predicted molecular mass of 23.8 kDa. **(B)** Cytosolic application of purified Lsm12 reduced lysosomal TPC2 currents activated by 0.3 µM PI(3,5)P_2_ in a concentration-dependent manner. For most Lsm12 concentrations, *n* = 10; *n* = 7 for 0.1 µM Lsm12 and *n* = 8 for the washout condition (“0 again”). **(C)** Purified Lsm12 (2 µM) markedly reduced endolysosomal TPC1 currents activated by 0.3 µM PI(3,5)P_2_ (*n* = 7). **(D)** Purified Lsm12 (2 µM) had little effect on endolysosomal TRPML1 currents activated by 0.3 µM PI(3,5)P_2_ (*n* = 6). Left panels in B–D: representative current-voltage (I-V) relationships obtained using a voltage ramp from −120 mV to +120 mV, together with schematic illustrations of whole-endolysosome recordings showing the major cations on either side of the membrane, from vacuolin-1-enlarged endolysosomes acutely isolated from HEK293 cells. Right panels in B–D: summary of current amplitudes measured at −120 mV (B, D) or +120 mV (C), showing individual data points and mean ± SEM, normalized to the corresponding current recorded in the absence of Lsm12. *, p < 0.05; **, p < 0.01; ***, p < 0.001; ****, p < 0.0001.

### Lsm12 competitively reduces TPC2 sensitivity to PI(3,5)P_2_

To investigate the mechanism underlying Lsm12-mediated inhibition of TPCs, we focused on TPC2 using the plasma membrane–targeted mutant TPC2-L11A/L12A (TPC2^PM^) (*22*). Inside-out patch-clamp recordings were performed in HEK293 cells transiently expressing TPC2^PM^. Perfusion of purified Lsm12 onto the cytosolic side of excised patches produced rapid (τ = 17.1 ± 1.4 s, n = 5) and near-complete inhibition of PI(3,5)P_2_-activated currents at 0.5 µM within ∼100 s (**Fig. 2A**). Upon washout of Lsm12 in the continued presence of PI(3,5)P_2_, the inhibited currents fully recovered (**Fig. 2A, B**). The recovery time constant was 11.5 ± 2.0 s (n = 5), not significantly different from the activation time constant measured with 300 nM PI(3,5)P_2_ in the absence of Lsm12 pretreatment (9.0 ± 1.5 s, n = 5) (**Fig. 2A, C**). Heat-inactivated Lsm12 at 0.5 or 5 µM showed little to no inhibition of currents activated by 0.3 or 1 µM PI(3,5)P_2_, respectively (**Fig. 2D**), conditions under which the active protein produced nearly complete inhibition (**Fig. 2B** and below), indicating that inhibition requires functional Lsm12.

**Figure 2.**
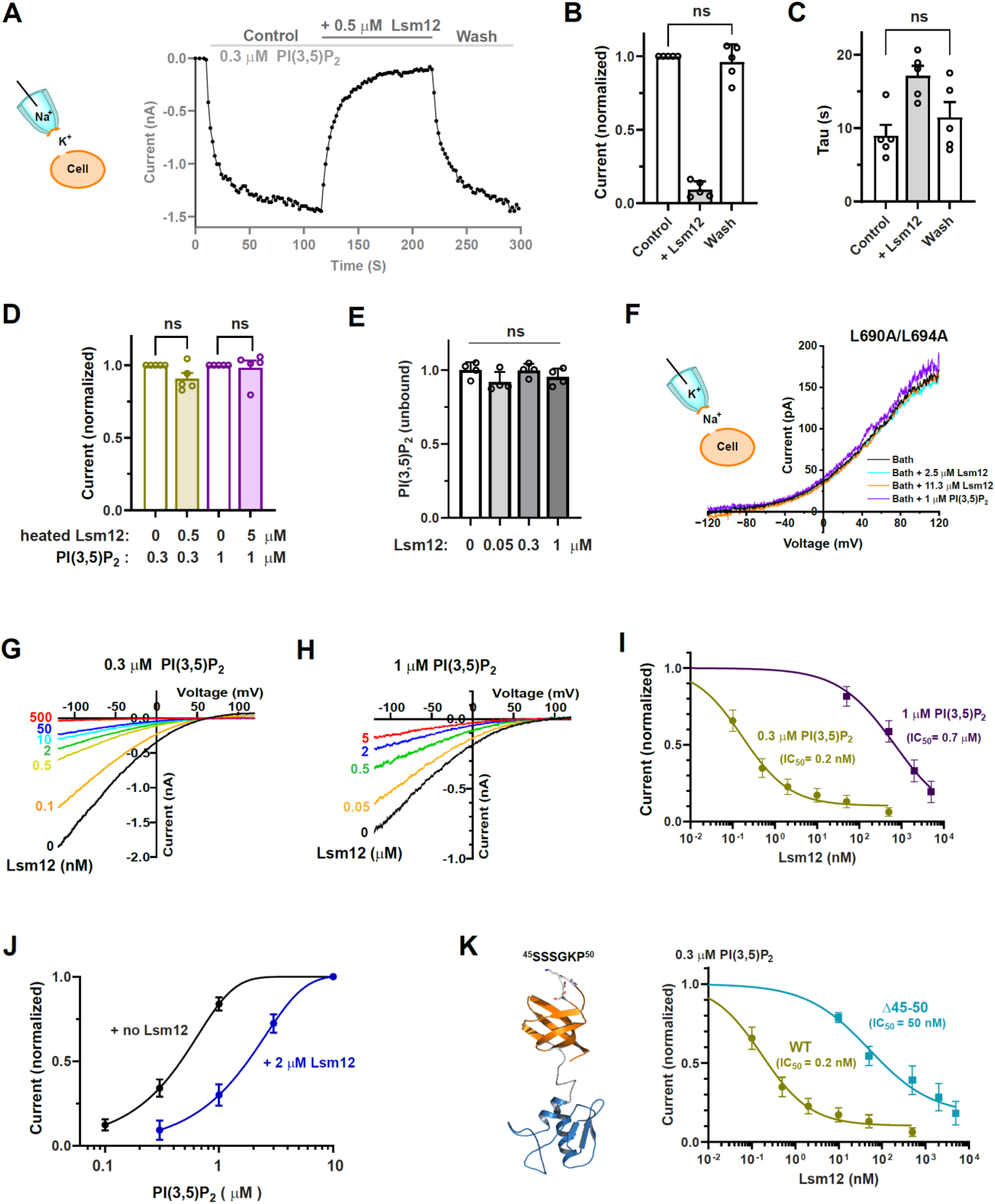
Lsm12 competitively inhibits PI(3,5)P_2_ activation of TPC2. **(A)** Representative time course of TPC2^PM^ currents at −120 mV during perfusion with 0.3 µM PI(3,5)P_2_. After current activation under control conditions, 0.5 µM Lsm12 was applied to inhibit the current, and subsequent washout of Lsm12 led to recovery. **(B, C)** Summary of the normalized current amplitudes derived from the maximal (activation) and minimal current (inhibition) levels (**B**) and time constants (τ) (**C**) during the control, Lsm12 wash-on, and wash-off phases from recordings as shown in **A** (*n* = 5). **(D)** Effects of heat-treated Lsm12 on PI(3,5)P_2_-activated TPC2^PM^ currents, normalized to currents recorded in the absence of heat-treated Lsm12. **(E)** Fraction of unbound BODIPY FL PI(3,5)P_2_ remaining after incubation of 0.3 µM lipid with 0.05, 0.3, or 1 µM 6×His-tagged Lsm12 (*n* = 4), determined from the fluorescence remaining in solution after precipitation of bead-bound Lsm12 with IMAC beads. **(F)** Representative I–V relationships of constitutively active TPC2^PM^ carrying the L690A/L694A mutation, recorded under bath conditions alone or after addition of the indicated concentrations of Lsm12 or PI(3,5)P_2_. **(G, H)** Representative I–V relationships of TPC2^PM^ currents activated by 0.3 µM (**G**) or 1 µM (**H**) PI(3,5)P_2_ in the absence and presence of the indicated concentrations of Lsm12. **(I)** Concentration-response relationships for inhibition by Lsm12 of TPC2^PM^ currents activated by 0.3 µM (*n* = 10) or 1 µM (*n* = 9) PI(3,5)P_2_. **(J)** PI(3,5)P_2_ concentration-response relationships for TPC2^PM^ currents in the absence and presence of 2 µM Lsm12. **(K)** Lsm12 concentration-response relationships for TPC2^PM^ pseudo-WT (*n* = 10) and Δ45–50 mutant (*n* = 8) channels activated by 0.3 µM PI(3,5)P_2_. Left, modeled structure of Lsm12 highlighting residues 45–50 in stick representation. Summarized data are presented as plots showing mean ± SEM, with or without individual data points, as indicated. ns, not significant (*P* > 0.05).

Lsm12 could inhibit TPC activation through sequestration of PI(3,5)P_2_ (reducing ligand availability), direct blockade of the channel pore, or modulation of channel sensitivity to PI(3,5)P_2_. To test PI(3,5)P_2_ sequestration, we measured unbound BODIPY FL PI(3,5)P_2_ after incubation with 6×His-tagged Lsm12 followed by IMAC bead pull-down. Under conditions in which most Lsm12 was precipitated, fluorescence of BODIPY FL PI(3,5)P_2_ remaining in solution was not significantly different from that with IMAC beads alone (**Fig. 2E**), indicating negligible binding between Lsm12 and PI(3,5)P_2_. Furthermore, Lsm12 had no effect on the constitutively active TPC2^PM^-L690A/L694A channels we previously identified (*40*) (**Fig. 2F**), ruling out pore blockade and indicating dependence on ligand-driven activation.

We next examined the relationship between Lsm12 inhibition and PI(3,5)P_2_ concentration. At 0.3 µM PI(3,5)P_2_ (near the EC_50_), Lsm12 inhibited TPC2^PM^ currents with high potency (IC_50_ ∼0.2 nM) (**Fig. 2G, I**). In contrast, at 1 µM PI(3,5)P_2_, ∼1000-fold higher Lsm12 concentrations were required (IC_50_ ∼0.7 µM) (**Fig. 2H, I**). For example, 2 and 50 nM Lsm12 inhibited 0.3 µM PI(3,5)P_2_–activated currents by 77% and 93%, respectively, whereas 2 and 5 µM Lsm12 produced 67% and 81% inhibition at 1 µM PI(3,5)P_2_. The pronounced shift in Lsm12 potency with increasing PI(3,5)P_2_ concentration indicates that Lsm12 and PI(3,5)P_2_ functionally oppose each other, consistent with a competitive mechanism. Accordingly, Lsm12 reduced the apparent sensitivity of TPC2^PM^ to PI(3,5)P_2_. The PI(3,5)P_2_ dose–response curve was right-shifted in the presence of 2 µM Lsm12, with the EC_50_ increasing from ∼0.4 µM to ∼2 µM (**Fig. 2J**). Deletion of residues 45–50 (a loop region of the Lsm domain) in Lsm12, which reduces interaction with TPC2 (*33*), markedly impaired inhibition, increasing the IC_50_ to ∼50 nM (∼250-fold shift) (**Fig. 2K**). Together, these data support a model in which Lsm12 competitively reduces TPC2 sensitivity to PI(3,5)P_2_ through interactions with TPC2.

### NAADP elicits TPC2^PM^ Na^+^ currents in the presence of PI(3,5)P_2_ by relieving inhibition from Lsm12

Because Lsm12 functions as an NAADP receptor (*33*), we examined how NAADP modulates Lsm12-mediated inhibition of PI(3,5)P_2_-activated TPC2. Excised plasma membrane patches from TPC2^PM^-expressing HEK293 cells were perfused with NAADP in the presence of 0.3 µM PI(3,5)P_2_ and 50 nM purified Lsm12, a condition in which PI(3,5)P_2_-evoked currents were largely suppressed (**Fig. 3A**). (**Fig. 3A**). Interestingly, addition of NAADP (10 nM–1 µM) restored currents in a dose-dependent manner, increasing from 13% (Lsm12 alone) to 36% at 100 nM and 69% at 1 µM (normalized to PI(3,5)P_2_ alone) (**Fig. 3A**). Consistent with the high specificity of Lsm12 for NAADP over NADP (*33, 41*), 1 µM NADP had no effect on TPC2^PM^ currents under the same conditions (**Fig. 3B**), indicating that current restoration requires specific NAADP–Lsm12 interactions.

**Figure 3.**
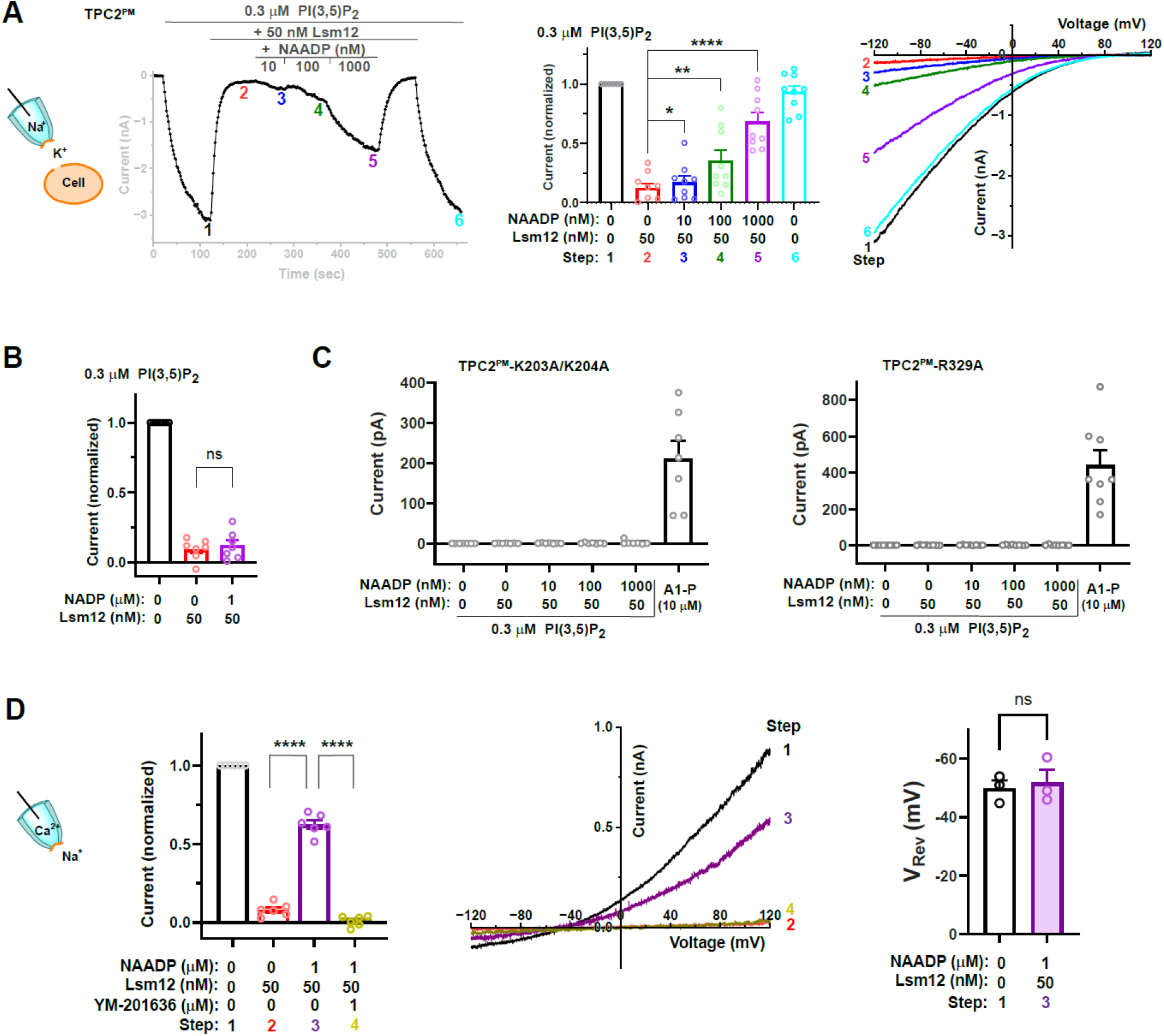
NAADP elicits TPC2^PM^ Na^+^ currents in the presence of PI(3,5)P_2_ by relieving inhibition from Lsm12. (**A**) NAADP elicits TPC2^PM^ currents in the presence of PI(3,5)P_2_ and Lsm12. **Left**: representative time course of TPC2^PM^ currents recorded at −120 mV during continuous application of 0.3 µM PI(3,5)P_2_, followed by sequential addition of 50 nM Lsm12 alone and then 50 nM Lsm12 together with increasing concentrations of NAADP (10, 100, and 1000 nM), and subsequent withdrawal of NAADP and Lsm12. **Middle**: summary of current amplitudes measured at −120 mV and normalized to the initial current activated by 0.3 µM PI(3,5)P_2_ alone. **Right**: representative I–V relationships obtained under the corresponding conditions. (**B**) NADP does not elicit TPC2^PM^ currents in the presence of PI(3,5)P_2_ and Lsm12. Current amplitudes were measured at −120 mV and normalized to the initial current activated by 0.3 µM PI(3,5)P_2_ alone. (**C**) NAADP fails to induce currents of PI(3,5)P_2_-insensitive mutant TPC2^PM^ channels. Current amplitudes were measured at −120 mV under the indicated conditions. Path-clamp recordings in (A-C) were performed under the same configuration and ionic conditions. **(D)** NAADP does not alter the relative Na^+^/Ca^2+^ permeability of PI(3,5)P_2_-activated TPC2^PM^ currents. **Middle:** representative I–V relationships under the indicated conditions. **Left and right:** summaries of normalized current amplitudes at +120 mV and reversal potentials, respectively. For reversal potential analysis, only I–V plots with maximal current amplitudes (+120 mV) between 0.5 and 1 nA were included to minimize error. Throughout this figure, the numbered steps and time points correspond to the color-coded conditions. Unless otherwise specified, the patch-clamp configurations and ionic conditions are illustrated schematically on the left. Bar graphs show individual data points and mean ± SEM. *, *p* < 0.05; **, p < 0.01; ****, *p* < 0.0001; ns, *p* > 0.05.

To test whether channel activation by PI(3,5)P_2_ is required for NAADP-evoked currents, we examined PI(3,5)P_2_-insensitive mutants, TPC2^PM^-R329A and TPC2^PM^-K203A/K204A (*8*). These mutants were unresponsive to PI(3,5)P_2_ but remained activatable by TPC2-A1-P (10 µM) (**Fig. 3C**). NAADP failed to elicit currents in the presence of PI(3,5)P_2_ and Lsm12 in either mutant (**Fig. 3C**), indicating that NAADP acts through PI(3,5)P_2_-dependent channel activation.

To determine whether NAADP-evoked, Lsm12- and PI(3,5)P_2_-dependent TPC2^PM^ currents differ in ion permeability from those activated by PI(3,5)P_2_ alone, we recorded TPC2^PM^ currents under a bi-ionic condition with extracellular Ca²⁺ and intracellular Na⁺ in an inside-out configuration (**Fig. 3D**). YM-201636 (*40*) abolished NAADP-evoked currents in the presence of PI(3,5)P_2_ and Lsm12, confirming their origin from TPC2^PM^ (Fig. 3D). (**Fig. 3D**). Reversal potentials under Ca²⁺/Na⁺ bi-ionic conditions were similar for currents evoked by PI(3,5)P_2_ alone (−50 mV) and by NAADP in the presence of PI(3,5)P_2_ and Lsm12 (−52 mV) (**Fig. 3D**), indicating unchanged Na⁺/Ca²⁺ permeability. Together, these data show that NAADP elicits TPC2^PM^ Na⁺ currents by relieving Lsm12-mediated inhibition of PI(3,5)P_2_-dependent channel activation.

### NAADP restores PI(3,5)P_2_-dependent endolysosomal TPC activities inhibited by Lsm12

To determine whether NAADP similarly regulates endolysosomal TPCs, we recorded whole-endolysosome currents of TPC2 and TPC1 under perfusion with PI(3,5)P_2_, purified Lsm12, and NAADP. Under conditions in which 1 µM Lsm12 suppressed currents evoked by 0.3 µM PI(3,5)P_2_, NAADP (100 nM and 1 µM) restored TPC2 currents in a dose-dependent manner, from 16% (Lsm12 alone) to 47% and 92% of the initial PI(3,5)P_2_-activated currents, respectively (**Fig. 4A**). In Lsm12-knockout (KO) cells, 1 µM NAADP co-applied with purified Lsm12 (2 µM) in the absence of PI(3,5)P_2_ elicited only minimal currents (25 ± 11 pA, n = 11) (**Fig. 4B, C**). In contrast, robust currents were observed when both Lsm12 (2 µM) and PI(3,5)P_2_ (0.3 µM) were present (490 ± 94 pA, n = 11) (**Fig. 4C**), indicating that NAADP-evoked currents require PI(3,5)P_2_-dependent channel activation, consistent with observations in TPC2^PM^ (**Fig. 3C**). Similar to TPC2, NAADP also relieved Lsm12-mediated inhibition of endolysosomal TPC1. In the presence of 1 µM Lsm12, 1 µM NAADP restored TPC1 currents from 22% to 80% of the currents activated by PI(3,5)P_2_ (0.3 µM) alone (**Fig. 4D**). Together, these results demonstrate that NAADP restores PI(3,5)P_2_-dependent endolysosomal TPC activity by relieving Lsm12-mediated inhibition, consistent with observations in TPC2^PM^ channels.

**Figure 4.**
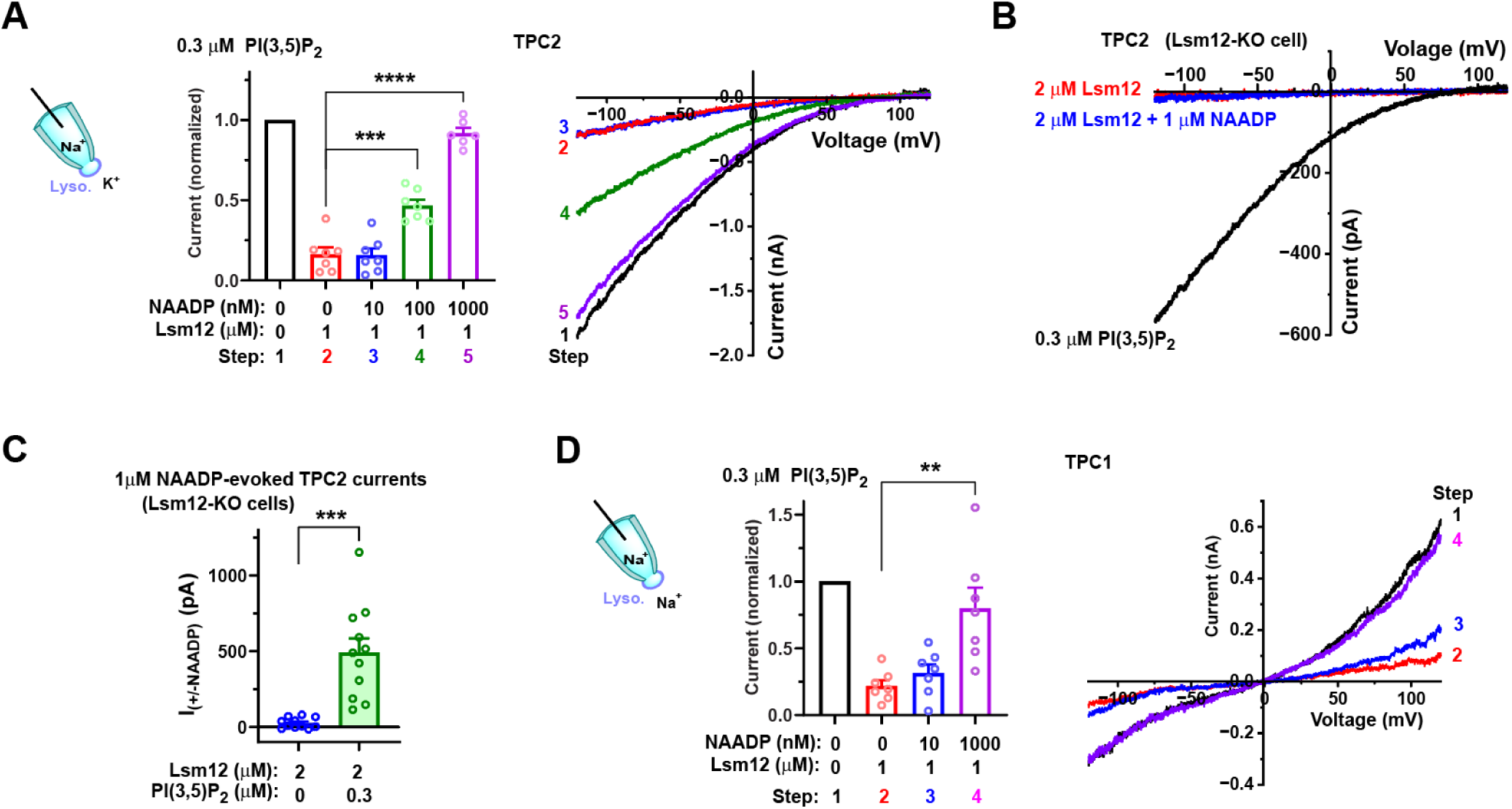
NAADP restores PI(3,5)P_2_-dependent endolysosomal TPC activities inhibited by Lsm12. **(A)** Summary of current amplitudes and representative I-V relationships showing that NAADP at 0.1 or 1 µM elicits currents of endolysosomal TPC2 in the presence of 0.3 µM PI(3,5)P_2_ and 1 µM Lsm12. (**B**) Representative I–V relationships comparing Lsm12 alone and Lsm12 plus NAADP, showing that NAADP produces minimal activation of TPC2 currents from Lsm12-KO cells in the absence of PI(3,5)P_2_. (**C**) Summary of 1 µM NAADP-induced currents in Lsm12-KO cells supplemented with purified Lsm12 (2 µM), in the presence or absence of 0.3 µM PI(3,5)P_2_. (**D**) Summary of current amplitudes and representative I-V relationships showing that NAADP (1 µM) elicits currents of endolysosomal TPC1 in the presence of 0.3 µM PI(3,5)P_2_ and 1 µM Lsm12. Current amplitudes for summary plots were measured at −120 mV in (A) and (C), and at +120 mV in (D). Where indicated, currents were normalized to those recorded in the absence of NAADP and Lsm12 within the same recording. Patch-clamp configurations and ionic conditions were identical for (A–C) and are illustrated schematically to the left of (A) and (D). Bar graphs show individual data points and mean ± SEM. **, p < 0.01; ***, *p* < 0.001; ****, *p* < 0.0001.

### Endogenous Lsm12 modulates TPC2 activation

Given the strong effect of purified Lsm12 on TPC gating, we asked whether endogenous Lsm12 exerts similar regulation. To test this, we examined TPC2^PM^ activity in the presence of the membrane-permeable agonist TPC2-A1-P, a functional analogue of PI(3,5)P_2_ (*42*), since PI(3,5)P_2_ cannot be directly applied intracellularly. In inside-out patches, 2 µM Lsm12 had little effect on TPC2^PM^ currents activated by 40 µM TPC2-A1-P, a near-maximal effect condition (*42*) (**Fig. 5A**). In contrast, when currents were elicited by 10 µM TPC2-A1-P, 2 µM Lsm12 inhibited ∼67% of the current (**Fig. 5B**). Addition of 1 µM NAADP did not relieve this inhibition (**Fig. 5B**), highlighting differences between PI(3,5)P_2_ and TPC2-A1-P in TPC2 activation, consistent with the ability of TPC2-A1-P to activate PI(3,5)P_2_-insensitive TPC2^PM^ mutants (**Fig. 3C**).

**Figure 5.**
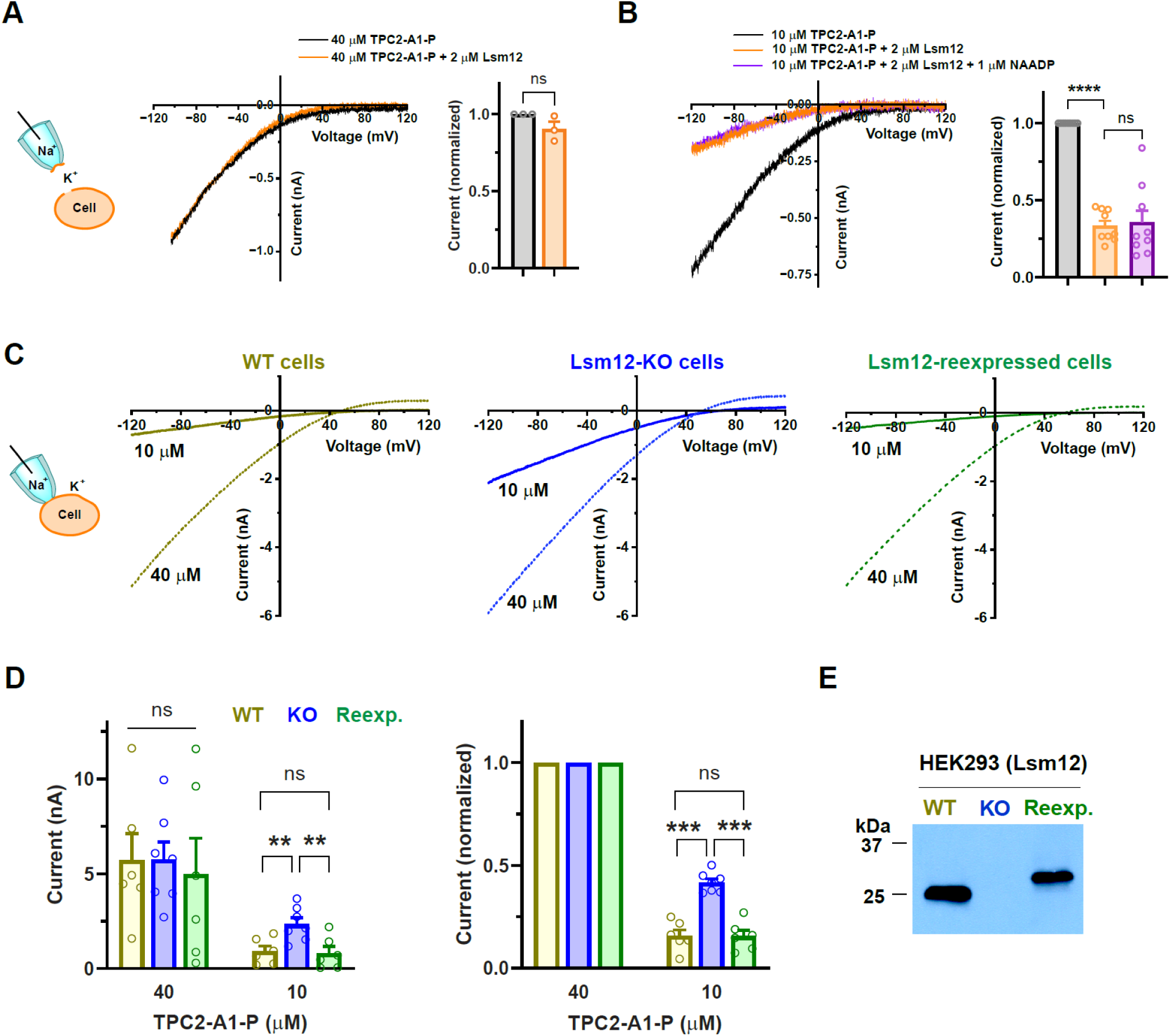
Endogenous Lsm12 inhibits TPC2. (**A, B**) Representative I–V relationships and corresponding summaries of TPC2^PM^ currents activated by 40 µM (A) or 10 µM (B) TPC2-A1-P. Currents were recorded in the inside-out excised patch configuration in the absence or presence of 2 µM Lsm12 or 1 µM NAADP, as indicated. For summary plots, current amplitudes were measured at −120 mV and normalized to those recorded in the absence of Lsm12. Conditions are color-coded as indicated. (**C**) Representative I–V relationships of TPC2^PM^ currents activated by extracellularly applied TPC2-A1-P at the indicated concentrations and recorded in the cell-attached configuration from WT, Lsm12-KO, and Lsm12-reexpressing HEK293 cells. (**D**) Summary of TPC2^PM^ current amplitudes measured at −120 mV, shown as absolute amplitudes (left) or normalized to currents elicited by 40 µM TPC2-A1-P in the same recording (right). (**E**) Immunoblot of lysates from WT, Lsm12-KO, and Lsm12-reexpressing HEK293 cells probed with an anti-Lsm12 antibody. For Lsm12 reexpression, a C-terminal FLAG- and 6×His-tagged construct was used. Cell types are color-coded as indicated in (C–E). Bar graphs show individual data points and mean ± SEM. *, *p* < 0.05; **, p < 0.01; ***, *p* < 0.001; ns, *p* > 0.05.

We next assessed the role of endogenous Lsm12 in intact cells using cell-attached recordings of TPC2^PM^ in WT, Lsm12-KO, and Lsm12-reexpressing HEK293 cells. Robust currents were elicited by extracellular application of TPC2-A1-P (**Fig. 5C, D**), consistent with its membrane permeability. At 40 µM TPC2-A1-P, current amplitudes were comparable across WT, KO, and reexpressing cells (**Fig. 5C, D**). However, at 10 µM, currents in Lsm12-KO cells were ∼2.6-fold larger than in WT cells (**Fig. 5C, D**). Reexpression of Lsm12 restored responses to levels similar to WT (**Fig. 5C, D**). This difference was more pronounced after normalization to 40 µM responses, reducing patch-to-patch variability (**Fig. 5D**). Immunoblotting with an anti-Lsm12 antibody confirmed loss and reexpression of Lsm12 in KO and rescued cells, respectively (**Fig. 5E**). Together, these results indicate that endogenous Lsm12 negatively regulates PI(3,5)P_2_-dependent TPC2 activity.

To test whether endogenous Lsm12 modulates lysosomal TPC2 activation, we measured TPC2-A1-P–induced intracellular Ca2+ signals in TPC2-expressing WT and Lsm12-KO cells using cell microinjection and GCaMP6-based Ca²⁺ imaging, as we previously described (*33, 40*). In both cell types, TPC2-A1-P (1–50 µM) elicited similar responses across concentrations in cells lacking exogenous TPC expression, likely reflecting TPC-independent effects (**Supplementary Fig. 1**). However, in cells expressing exogenous TPC2, TPC2-A1-P (≥10 µM) elicited additional TPC2-dependent Ca²⁺ elevations, exhibiting a bell-shaped dose–response (**Supplementary Fig. 1**), resembling that of NAADP (*43*). Within the tested range (1, 10, 50, and 100 µM), Lsm12-KO cells displayed greater sensitivity to TPC2-A1-P than WT cells. Specifically, Lsm12-KO cells showed maximal responses at 10 µM, whereas WT cells responded weakly at 10 µM and reached maximal responses at 50 µM (**Supplementary Fig. 1**). These results indicate that endogenous Lsm12 suppresses lysosomal TPC2 activity.

### Endogenous PI(3,5)P_2_ contributes to NAADP-evoked Ca^2+^ release

TPCs mediate NAADP-evoked Ca²⁺ release from acidic endolysosomal stores. As PI(3,5)P_2_ is the only known endogenous direct activator of TPCs, we tested whether it is required for NAADP-evoked Ca²⁺ release. Acute depletion of endogenous PI(3,5)P_2_, achieved by co-injecting NAADP with either a PI(3,5)P_2_ antibody or a PI(3,5)P_2_-binding GRIP protein, significantly reduced NAADP-evoked increases in Ca²⁺ indicator (GCaMP) fluorescence by 57% and 50%, respectively, compared with control PI(4,5)P2 antibody or GRIP protein (**Fig. 6A**). These results suggest that endogenous PI(3,5)P_2_ contributes acutely to NAADP-evoked Ca²⁺ release. Although the precise role of PI(3,5)P_2_ in NAADP signaling remains to be defined, our findings that NAADP restores TPC activity by relieving Lsm12-mediated inhibition of PI(3,5)P_2_-activated channels (**Fig. 6B**), providing a Lsm12-mediated mechanistic link between NAADP signaling and PI(3,5)P_2_-dependent TPC activation.

**Figure 6.**
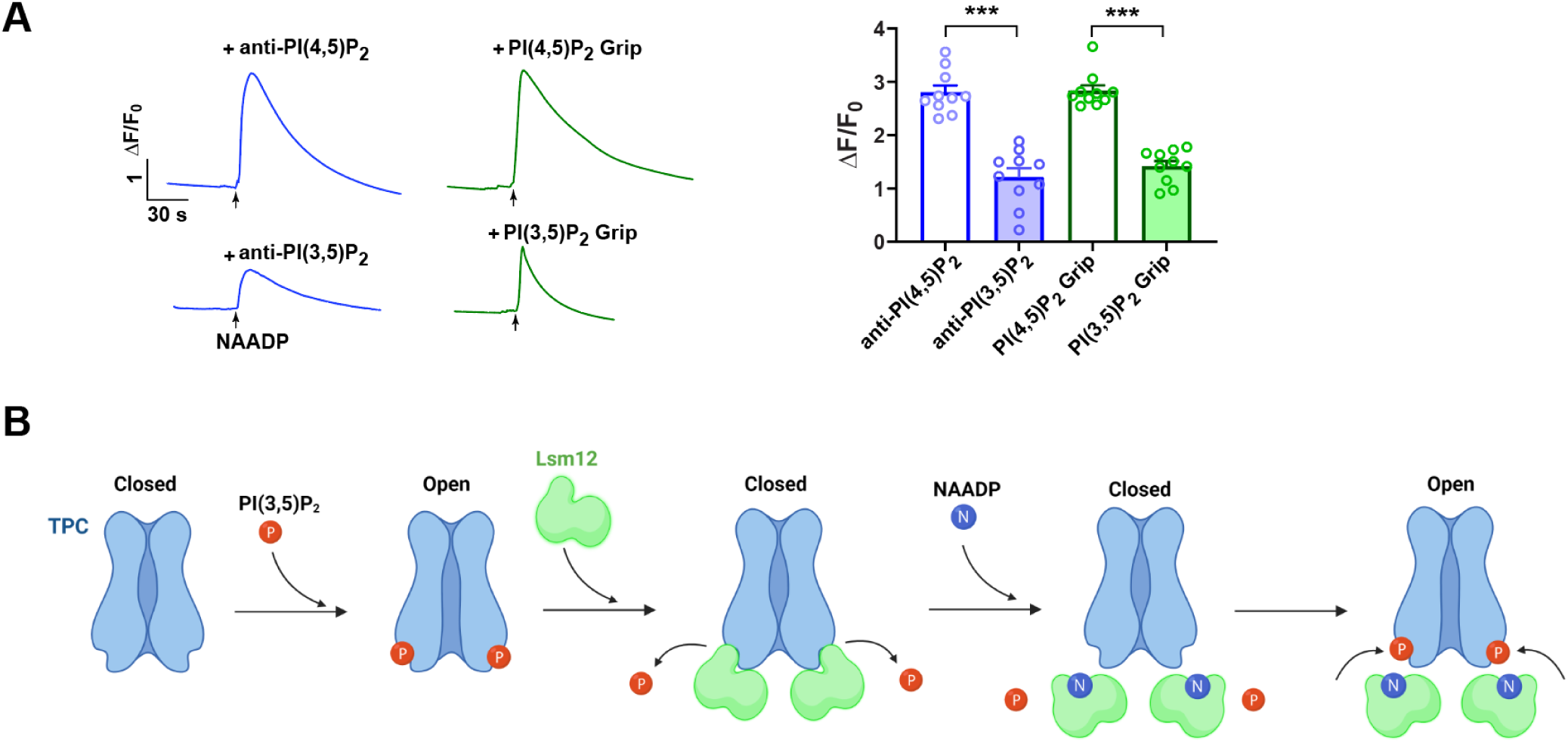
Interplay between NAADP and PI(3,5)P_2_ in TPC regulation and NAADP-evoked Ca²⁺ release. **(A)** Endogenous PI(3,5)P_2_ contributes to NAADP-evoked Ca²⁺ release. Antibodies or lipid-binding “grip” proteins targeting PI(3,5)P_2_ or PI(4,5)P2 were co-injected with NAADP into cells. Left: Representative Ca²⁺ indicator traces. Right: Summary of peak NAADP-evoked Ca²⁺ responses. Bar graphs show individual data points and mean ± SEM. ***, *p* < 0.001. (**B**) Schematic summary of the interplay among PI(3,5)P_2_, Lsm12, and NAADP in TPC regulation. PI(3,5)P_2_ activates TPCs, whereas Lsm12 binding to TPCs competitively suppresses PI(3,5)P_2_-activated channels. NAADP binding to Lsm12 relieves this suppression, restoring channel opening in the presence of PI(3,5)P_2_. The interacting sites of the ligand and proteins, as well as the number of Lsm12 molecules depicted in the cartoon, are arbitrary.

## DISCUSSION

The recent identification of NAADP receptor proteins supports a unifying view in which NAADP evokes Ca²⁺ release from endolysosomal acidic stores via receptor proteins and their associated ion channels, particularly TPCs. However, the molecular mechanism by which NAADP, through its receptors, is coupled to TPC activation remains unclear. In this study, we investigated the functional interplay among three TPC regulators: PI(3,5)P_2_, NAADP, and the NAADP receptor Lsm12. Our results reveal a model in which Lsm12 competitively inhibits PI(3,5)P_2_ activation of TPCs, and NAADP does not directly activate TPCs but instead relieves Lsm12-mediated inhibition of PI(3,5)P₂-dependent gating (**Fig. 6B**). This model provides a framework for reconciling two prominent observations in the literature: (i) TPCs are implicated in NAADP evokes Ca²⁺ release from acidic stores in many experimental systems; (ii) the lack of a defined mechanism for NAADP action on TPCs, and even the lack of consensus as to whether NAADP actually gates TPCs. In this context, NAADP and PI(3,5)P_2_ do not function independently but, through Lsm12, operate as an integrated signaling module in which channel activity reflects the balance among lipid-dependent activation, Lsm12-mediated inhibition, and NAADP-induced relief of inhibition.

### Lsm12 as a competitive modulator of PI(3,5)P_2_ gating

Our data strongly support a model in which Lsm12 acts as a competitive negative regulator of PI(3,5)P_2_-dependent TPC activation. Several observations converge on this conclusion. First, Lsm12 inhibition is highly dependent on PI(3,5)P_2_ concentration and produces a rightward shift in the PI(3,5)P_2_ dose–response relationship. Second, Lsm12 does not sequester PI(3,5)P_2_ and does not inhibit constitutively active TPC mutants, arguing against nonspecific lipid buffering or pore blockade. Third, weakening of the Lsm12–TPC interaction markedly reduces inhibition, indicating that direct interaction with the channel is required. Fourth, the Lsm12–TPC2 interaction appears to be dynamic, as the inhibition is rapidly reversible, at least in the presence of PI(3,5)P_2_. Together, these findings are most consistent with a mechanism of competitive inhibition in which Lsm12 binding to TPCs reduces channel sensitivity to PI(3,5)P_2_, whereas PI(3,5)P_2_ binding reciprocally weakens the Lsm12–TPC interaction. This mechanism is distinct from that of a pore blocker or a typical noncompetitive inhibitor, which acts largely independently of ligand concentration by occluding ion permeation or impairing coupling between ligand binding and channel opening, and primarily reduce maximal current amplitude.

### Lsm12 as a tonic suppressor of TPC activity

Our data further suggest that endogenous Lsm12 likely exerts tonic suppression of TPC activity under basal conditions. Genetic loss of Lsm12 enhances TPC2 responsiveness to TPC2-A1-P, whereas re-expression restores inhibition. These findings indicate that Lsm12 is not merely an accessory factor required for NAADP signaling, but rather functions as an active negative regulator of channel gating by ambient PI(3,5)P_2_. This tonic inhibition is likely physiologically important in the context of the lipid environment of endolysosomal membranes. Although PI(3,5)P_2_ is a low-abundance phosphoinositide, its enrichment on endolysosomal membranes is expected to reach locally sufficient levels to support basal activity of PI(3,5)P_2_-sensitive channels. In this context, Lsm12-mediated inhibition would restrain constitutive channel opening, thereby preventing excessive cation flux across endolysosomal membranes and supporting organelle homeostasis. Unchecked TPC activity could lead to dysregulated luminal ion composition, altered membrane potential, and defects in endolysosomal trafficking or fusion. Thus, the broadly expressed Lsm12 likely serves as a tonic “brake” that sets the basal activity of TPCs, ensuring that channel activation is tightly coupled to physiological stimuli rather than the ambient presence of PI(3,5)P_2_.

### NAADP relieves Lsm12 inhibition of PI(3,5)P_2_-dependent TPC activation without directly activating the channels

In our recording conditions, NAADP even in the presence of supplemental Lsm12 failed to induce significant TPC2 currents. However, NAADP elicits robust large PI(3,5)P_2_-dependent TPC currents in the presence of Lsm12 by lifting the Lsm12-mediated inhibition. The requirement for PI(3,5)P_2_, the inability of NAADP to activate PI(3,5)P_2_-insensitive mutants, and the lack of effect of NADP strongly support this interpretation. Within this framework, NAADP functions not directly as a channel agonist but as a relief signal that counteracts Lsm12-mediated inhibition. Thus, the NAADP-dependent activation of TPCs observed in this study is conditional upon pre-existing lipid-dependent gating machinery. Such a mechanism allows cells to maintain tight control over basal TPC activity from activation by PI(3,5)P_2_ while enabling rapid activation in response to NAADP.

The findings from this study provide an explanation for the difficulty or variability in detecting NAADP-evoked TPC currents. If NAADP acts by relieving inhibition rather than directly activating the channel, then measurable responses require sufficient PI(3,5)P_2_ availability and presence of functional Lsm12. Recording conditions such as excised plasma membrane patches or enlarged endolysosomes are expected to abolish or attenuate NAADP responses, due to the loss of cytosolic Lsm12 components or depletion of PI(3,5)P_2_ following vacuolin-1 treatment (*44*). Our findings therefore underscore the importance of cellular context and lipid signaling in NAADP-dependent TPC activation.

In our previous work (*33*), NAADP-evoked currents were observed when NAADP was microinjected into cells during whole-cell patch-clamp recordings of TPC2^PM^. It remains to be determined whether the observed currents are PI(3,5)P_2_-dependent as PI(3,5)P_2_ is also present on plasma membranes although not enriched (*45*). Future biochemical investigations will be valuable to further delineate whether Lsm12 modulates PI(3,5)P_2_ binding to TPCs and how NAADP influences Lsm12–TPC physical interactions. Although prior co-pulldown experiments did not reveal a clear effect of NAADP on Lsm12–TPC2 association (*33*), such conventional co-immunoprecipitation approaches, particularly with stringent washing conditions, is unlikely to capture the dynamic and transient nature of these interactions suggested by our results.

### Coupling NAADP signaling to endolysosomal lipid state

Prior studies suggest a potential interplay between PI(3,5)P_2_ and NAADP in TPC regulation and NAADP-mediated Ca²⁺ signaling. This includes synergistic effects of NAADP and PI(3,5)P_2_ on TPC currents (*46, 47*), as well as the loss of NAADP-evoked Ca²⁺ release in PI(3,5)P_2_-insensitive TPC1 mutants (*48*). Our data further demonstrate that acute depletion of PI(3,5)P_2_, achieved by microinjection of antibodies and GRIP proteins, rapidly and significantly impairs NAADP-induced Ca²⁺ signals. This result agrees with our finding of the PI(3,5)P_2_-dependence in NAADP-induced TPC2 currents in the presence of Lsm12. Thus, PI(3,5)P_2_ is functionally important in NAADP signaling. This finding highlights a key principle: NAADP signaling is intrinsically coupled to the phosphoinositide composition of endolysosomal membranes. Because PI(3,5)P_2_ levels are dynamically regulated by lipid kinases and phosphatases and vary across endosomal maturation states, TPC activity, and thus NAADP signaling may be tightly linked to organelle identity, trafficking, and metabolic status.

### Contribution of Na⁺-selective TPC currents to Ca²⁺ signaling

PI(3,5)P_2_-activated TPCs are highly selective for Na⁺, with only minimal permeability to Ca²⁺. Our data indicate that NAADP-evoked TPC currents retain the ion selectivity characteristic of PI(3,5)P_2_-activated channels. The extent to which Na⁺-selective TPC currents contribute to NAADP-evoked Ca²⁺ signaling remains to be determined. Nevertheless, TPC2-A1-P, which functionally mimics PI(3,5)P_2_ in terms of TPC2 ion permeability, is capable of eliciting TPC-dependent intracellular Ca²⁺ elevation, as demonstrated in this study and previous reports (*42, 46*). Two non-mutually exclusive mechanisms may underlie this effect. First, despite the low Ca²⁺ permeability, the small Ca²⁺ influx through the Na^+^-selective TPCs in endolysosomes may be sufficient to trigger Ca²⁺-induced Ca²⁺ release from the endoplasmic reticulum via IP₃ or ryanodine receptors. Second, Na⁺ flux through TPCs may alter lysosomal membrane potential or ionic gradients, thereby indirectly facilitating Ca²⁺ release through other channels or transporters.

### Conclusions and perspectives

Our work expands the regulatory landscape of TPCs and provides a mechanism of NAADP-evoked TPC activation. We found that TPC activity is governed by an integrated regulatory module in which PI(3,5)P_2_ provides positive gating input, Lsm12 imposes inhibition, and NAADP dynamically relieves this inhibition. This layered control mechanism may allow TPCs to function as coincidence detectors, integrating metabolic, lipid, and second messenger signals at endolysosomal membranes. Future biochemical and structural studies of the NAADP–Lsm12–TPC–PI(3,5)P_2_ axis will be needed to refine the molecular mechanisms underlying NAADP-mediated relief of Lsm12 inhibition of PI(3,5)P_2_-dependent activation. Furthermore, given reports of NAADP-evoked Ca^2+^ currents (*3, 39, 42, 46*), it will be important to determine whether such observations relate to the Lsm12-mediated, PI(3,5)P_2_-dependent mechanism identified here, and whether additional mechanisms or pathways may contribute to enhanced Ca^2+^ conductance.

## METHODS AND MATERIALS

### Bacterial expression and identification of Lsm12

The bacterial expression plasmid pET-6His-TB-Lsm12, which encodes human Lsm12 protein with an N-terminally tagged cleavable thrombin (TB) cleavage motif after 6×His tag, was constructed with bacterial pET vector and transformed into E. coli strain BL21 (DE3), as we previous report(*33*). The cells were grown at 37 °C and collected 4 h after induction with 1 mM IPTG. Cells were harvested by centrifugation and broken by sonication in 50 mM Tris-HCl pH 8.0, 500 mM NaCl, and 5% glycerol supplemented with 1 mM PMSF and PierceTM Protease inhibitor tablet (Cat# A32965 from ThermoFisher). After centrifugation at 16,000 × g for 40 min, the soluble fraction was collected as supernatant. The 6×His-tagged Lsm12 was purified by passing the soluble fraction through a Ni-NTA column connected with Biorad NGC chromatography system. After binding, Ni-NTA column was washed with gradient of imidazole from 10 mM to 200 mM and then eluted from the column with 300 mM imidazole in the same buffer. Imidazole and excessive salts in elute were removed by dialysis in 25 mM Tris-HCl pH 8.0 and 50 mM NaCl. The purified Lsm12 protein was identified by SDS-PAGE. Endogenous Lsm12 in HEK293 cells was checked by immunoblotting using rabbit monoclonal anti-Lsm12 antibody (Cat# EPR12282 from Abcam) at a dilution factor of 1:1000.

### Cell culture, plasmids, and transfection

HEK293 cells were cultured in Dulbecco’s modified Eagle’s medium with 10% fetal bovine serum, 1% penicillin, and streptomycin in a 5% CO_2_ incubator. Lsm12 knockout cell was generated and used as we reported before(*33*). Similar to our recent reports (*33*), recombinant cDNA constructs of human TPC1 (GenBank: AY083666.1) and human TPC2 (GenBank: BC063008.1) with FLAG and V5 epitopes on their C-termini were constructed with pCDNA6 vector (Invitrogen). To facilitate identification of transfected cells, an IRES-containing bicistronic vector, pCDNA6-TPC2-V5-IRES-AcGFP (*33*), was used in the electrophysiological experiments. Mutations were made with QuikChange II XL Site-Directed Mutagenesis Kit (Agilent Technologies). Cells were transiently transfected with plasmids with transfection reagent of PolyJet (SignaGen Laboratories), Lipofectamine 2000 (Invitrogen), or polyethylenimine “Max” (PEI Max from Polysciences) and subjected to experiments within 16-48 h after transfection. For the cell health of the mutant L690A/694A after transfection, 2 µM YM201636 was added into the complete serum medium to block TPC2. For human TPC1 channels, pEGFP-C1 was cotransfected at the same time to identify transfected cells for patch clamp recording. Cells were treated with 1% trypsin 4-6 h after transfection and seeded on polylysine-treated glass coverslips soaked in an incubator until recording.

### Electrophysiology

For inside-out and cell-attached patch-clamp recording, HEK293 cells were transiently transfected with plasma membrane–targeted human TPC2^L11A/L12A^ (TPC2^PM^). After 24 h of transfection, the channel currents were acquired at room temperature. For most inside-out recordings of excised plasma membrane patches and for cell-attached recordings of intact cells, the bath solution contained 145mM KMeSO_3_, 5mM KCl, and 10mM HEPES (pH 7.4), and the pipette solution contained 145mM NaMeSO_3_, 5mM NaCl, and 10mM HEPES (pH 7.4). For the constitutively open TPC2^PM^-L690A/694A mutant, the solutions were switched, i.e., the K^+^-based solution was used as the pipette solution, and the Na^+^-based solution was used in the bath instead. In some inside-out recordings, the bath solution was the Na^+^-based solution and the pipette solution was a Ca^2+^-based solution containing 105 mM CaCl2, 5mM HEPES, and 5 mM MES (pH 7.2) (pH was adjusted with Ca(OH)_2_). Whole lysosome patch-clamp recording of PI(3,5)P_2_-activated TPC2 activation was performed as previously reported by others and us. Cells were transfected with human TPC2 or human TPC1 plasmids using PolyJet for 4-6 hours and then treated with vacuolin-1 (1 μM) overnight to enlarge endolysosomes. For recordings of TPC2 currents, the cytoplasmic (bath) solution contained 145mM KMeSO_3_, 4mM NaCl 4, 0.39mM CaCl_2_, 1mM EGTA, and 10mM HEPES (pH 7.2) (pH was adjusted with KOH). The luminal (pipette) solution contained 140 mM NaMeSO_3_, 5mM KMeSO_3_, 2mM Ca(MeSO3)_2_, 1mM CaCl_2_, 10mM HEPES (pH 7.2) (pH was adjusted with NaOH). For recording TPC1 currents, the same Na^+^-based solution (pH 7.2) was used on both sides (pipette and bath). The TPC2 or TPC1 channel currents were elicited by perfusion of PtdIns(3,5)P2 diC8 (Echelon Biosciences #P-3058), TPC2-A1-P (MedChemExpress #HY-131615 or Sigma-Aldrich #SML3700), or TPC2-A1-N (MedChemExpress #HY-131614 or Sigma-Aldrich #SML3562) on the cytosolic side in inside-out recordings and whole endolysosomal recordings, or of TPC2-A1-P on the extracellular side in cell-attached recordings. Purified LSM12, NAADP (Tocris # 3905), or NADP (Sigma-Aldrich # 481972) were applied on the cytosolic side by perfusion. For all recordings, currents were measured using a voltage ramp protocol from −120 mV to +120 mV over 400 ms, applied every 2 s, with a holding potential of 0 mV. All data, except those obtained in the cell-attached mode, were recorded at a sampling rate of 10 kHz and low-pass filtered at 10 kHz using a MultiClamp 700B amplifier with pCLAMP software (Axon Instruments, Molecular Devices). In cell-attached mode, data were sampled at 50 kHz and filtered at 2.9 kHz using an EPC10 USB amplifier with PATCHMASTER software (HEKA Elektronik). Patch pipettes were polished to a resistance of 1.5–2 MΩ for inside-out and cell-attached recordings, and 5–8 MΩ for whole endolysosomal recordings. Dose curves were fitted by the Hill logistic equation. The time constant (τ) were acquired from singe exponential fitting.

### Evaluation of the binding between Lsm12 and PI(3,5)P_2_

500 µl 0.3 µM BODIPY FL PI(3,5)P_2_ (Echelon Biosciences #C-35F6) was incubated with designated concentrations (0, 0.05, 0.3, and 1 µM) of 6×His tagged Lsm12 for 10 min in the K^+^-based solution (145mM KMeSO3, 5mM KCl, and 10mM HEPES (pH 7.4)) that was used for inside-out patch-clamp recordings. 20 µl Nuvia IMAC (immobilized metal affinity chromatography) Ni-charged resin (Bio-Rad #7800800) pre-equilibrated with the K^+^-based solution was added and incubated for 10 min. After centrifugation to pellet the resin, the fluorescence of the unbound BODIPY FL PI(3,5)P_2_ in the supernatant was measured by a plate reader. The fractions of the unbound BODIPY FL PI(3,5)P_2_ were calculated from normalization of the measured fluorescence to that of the control without Lsm12. The efficiency of the Lsm12 pull-down by IMAC at similar conditions was estimated by using 6×His-tagged sfGFP-Lsm12 based on the loss of GFP fluorescence in solution after precipitation with IMAC resin.

### Imaging analysis of NAADP-evoked Ca^2+^ release

Ca^2+^ imaging analysis of NAADP-evoked Ca^2+^ elevation was performed as we recently described (*33*). Briefly, cells were co-transfected with cDNA constructs of human TPC2 and the Ca^2+^ reporter GCaMP6f, and the transfected cells were identified by GCaMP6f fluorescence. Cell microinjection was performed with a FemtoJet microinjector (Eppendorf). The pipette solution contained 110 mM KCl, 10 mM NaCl, and 20 mM Hepes (pH 7.2) supplemented with Dextran (10,000 MW)-Texas Red (0.3 mg/ml) and NAADP (100 nM), TPC2-A1-P (1, 10, 50 or 100 µM), or vehicle. When used, the PI(3,5)P_2_ and PI(4,5)P2 antibodies (Echelon Biosciences #Z-P035 and #Z-P045,) or Grip proteins (Echelon Biosciences #G-3501 and #G-4501) was added to the injection pipette solution at a final concentration of 5 µg/ml or 10 µg/ml (only for PI(3,5)P_2_ antibody). The bath was Hank’s balanced salt solution, which contained 137 mM NaCl, 5.4 mM KCl, 0.25 mM Na_2_HPO_4_, 0.44 mM KH_2_PO_4_, 1 mM MgSO_4_, 1 mM MgCl_2_, 10 mM glucose, and 10 mM HEPES (pH 7.4). To minimize interference by contaminated Ca^2+^, the pipette solution was treated with Chelex 100 resin (Sigma-Aldrich #C709) immediately before use. Microinjection (0.5 s at 150 hPa) was made ∼30 s after pipette tip insertion into cells. Only cells that showed no response to mechanical puncture, i.e., no change in GCaMP6f fluorescence for ∼30 s, were chosen for pipette solution injection. Successful injection was verified by fluorescence of the co-injected Texas Red. Fluorescence of GCaMP6f was monitored with an Axio Observer A1 microscope equipped with an AxioCam MRm digital camera and ZEN Blue 2 software containing a physiology module (Carl Zeiss) at a sampling frequency of 2 Hz. Elevation in intracellular Ca^2+^ concentration was reported by a change in fluorescence intensity measured as ΔF/F_0_, calculated from NAADP microinjection-induced maximal changes in fluorescence (ΔF at the peak) divided by the fluorescence immediately before microinjection (F_0_).

### Data analysis

The data was processed and plotted with Igor Pro (v5 or v6), GraphPad Prism (v10), or OriginLab (v2017 or 2020). All statistical values are performed as means + or ± SEM of *n* repeats of the experiments. All repeats in electrophysiological recordings or Ca^2+^ imaging analyses were obtained with distinct cells. Paired or unpaired Student’s t-test was used to calculate *p* values.

## ACKNOWLEDGMENTS

This work was supported in part by National Institutes of Health grants GM130814 (to J. Yan).

## AUTHOR CONTRIBUTIONS

X. Guan, C. Du, K. Shah, and J. Yan designed experiments, analyzed data, and wrote the manuscript. X. Guan, C. Du, and K. Shah performed experiments.

---

**Supplementary Figure 1.**
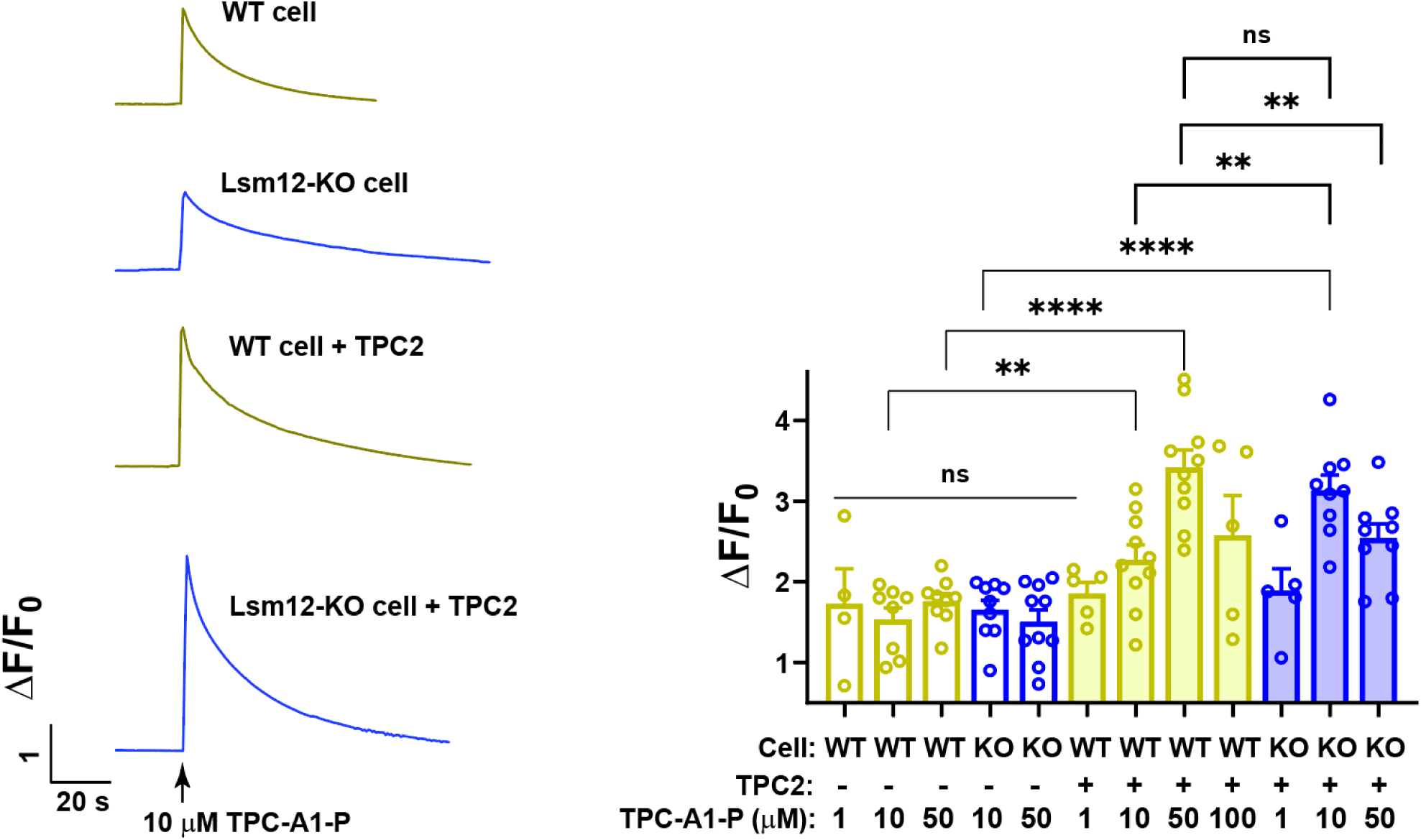
Lsm12 knockout enhances sensitivity of TPC2-expressing cells to TPC2-A1-P–induced Ca^2+^ elevation Left: Representative time courses of Ca^2+^ indicator signals in response to 10 µM TPC2-A1-P in WT and Lsm12-KO HEK293 cells, with or without TPC2 expression. **Right**: Summary of peak Ca^2+^ responses (ΔF/F₀) evoked by the indicated concentrations of TPC2-A1-P in WT and Lsm12-KO HEK293 cells, with or without TPC2 expression. Bar graphs show individual data points and mean ± SEM. **, *p* < 0.01; ***, *p* < 0.001; ****, *p* < 0.0001; ns, *p* > 0.05.

